# Pathogenic and Indigenous Denitrifying Bacteria are Transcriptionally Active and Key Multi-Antibiotic Resistant Players in Wastewater Treatment Plants

**DOI:** 10.1101/2020.12.14.422623

**Authors:** Ling Yuan, Yubo Wang, Lu Zhang, Alejandro Palomo, Jizhong Zhou, Barth F. Smets, Helmut Bürgmann, Feng Ju

## Abstract

The global rise and spread of antibiotic resistance greatly challenge the treatment of bacterial infections. Wastewater treatment plants (WWTPs) harbor and discharge antibiotic resistance genes (ARGs) as environmental contaminants. However, the knowledge gap on the host identity, activity and functionality of ARGs limits transmission and health risk assessment of WWTPs resistome. Hereby, a genome-centric quantitative metatranscriptomic approach was exploited to realize high-resolution qualitative and quantitative analyses of bacterial hosts of ARGs (i.e., multi-resistance, pathogenicity, activity and niches) throughout 12 urban WWTPs. We found that ∼45% of 248 recovered genomes expressed ARGs against multiple classes of antibiotics, among which bacitracin and aminoglycoside resistance genes in *Proteobacteria* was the most prevalent scenario. Both potential pathogens and indigenous denitrifying bacteria were transcriptionally active hosts of ARGs. The almost unchanged relative expression levels of ARGs in the most resistant populations (66.9%) and the surviving ARG hosts including globally emerging pathogens (e.g., *Aliarcobacter cryaerophilus*) in treated WWTP effluent prioritizes future examination on the health risks related with resistance propagation and human exposure in the receiving environment.

## INTRODUCTION

The extensive use of antibiotics and the resulting accelerated bacterial resistance dissemination have largely promoted the rise of antibiotic resistance as one of the greatest global public health threats^1, 2^. Most of the antibiotic wastes together with antibiotic resistant bacteria and antibiotic resistance genes (ARGs) emitted from anthropogenic sources in urban areas eventually enter wastewater treatment plants (WWTPs) which are considered as hotspots for the release of ARGs and their hosts into the environment^3–5^. The prevalence and high diversity of ARGs in WWTPs have been widely noted^4, 6–8^ through metagenomic approaches^9, 10^. However, the fragmented nature of reported metagenomic assemblies cannot solidly predict identity of ARG host. Previous study based on genome-centric metagenomics enables a better understanding of ARG hosts in activated sludge at the genome level^11^, but the lack of activity-based resistome monitoring make it impossible to examine expression activity of ARG and identify active ARG hosts in WWTPs.

Theoretically, genome-centric metatranscriptomics can overcome the above technical bottlenecks by providing both high-resolution genome-level taxonomy and global gene expression activities of environmental microorganisms. The host identity and activity of ARGs in activated sludge has been preliminarily explored with a genome-centric metatranscriptomic method^12^, but the absolute gene expression pattern of ARG hosts in the varying WWTP compartments (e.g., influent, activated sludge, effluent) remain unknown, restraining objective evaluation of environmental transmission and health risks of antibiotic resistance in the receiving environment of WWTPs effluent. Moreover, the important functional traits (e.g., nitrogen and phosphorus removal, pathogenicity and niche breadth) of ARG hosts are still poorly understood. The knowledge is, however, of particular interest as the locally-adapted microbes dedicated to organic and nutrients removal in activated sludge are under continuous and long-term exposure to subinhibitory levels of antimicrobial contaminants (e.g., antibiotics, heavy metals and biocides^6, 13, 14^). Considering the fact that enteric microbes including pathogens are being continuously introduced into WWTPs with sewage inflow, their regular close contact with indigenous microbes that are potentially under stress from exposure to antimicrobials may create conditions where resistance exchange involving pathogens followed by multi-resistance selection and potential local niche adaptation is favored (Fig. 1). This may represent expectable but not yet evaluated ecological and health risks^15^. Although functional bacteria^16–18^ and ARGs^6, 19, 20^ in WWTPs were extensively studied independently through culture-independent approaches^21–23^, the extent to which indigenous microbes and especially the key functional bacteria in different compartments of WWTPs may represent hitherto-unrecognized recipients or even disseminators of ARGs remains unexplored. Efforts are needed to fill all these knowledge gaps on the antibiotic resistance in WWTPs with improved methodology.

**Fig. 1.**
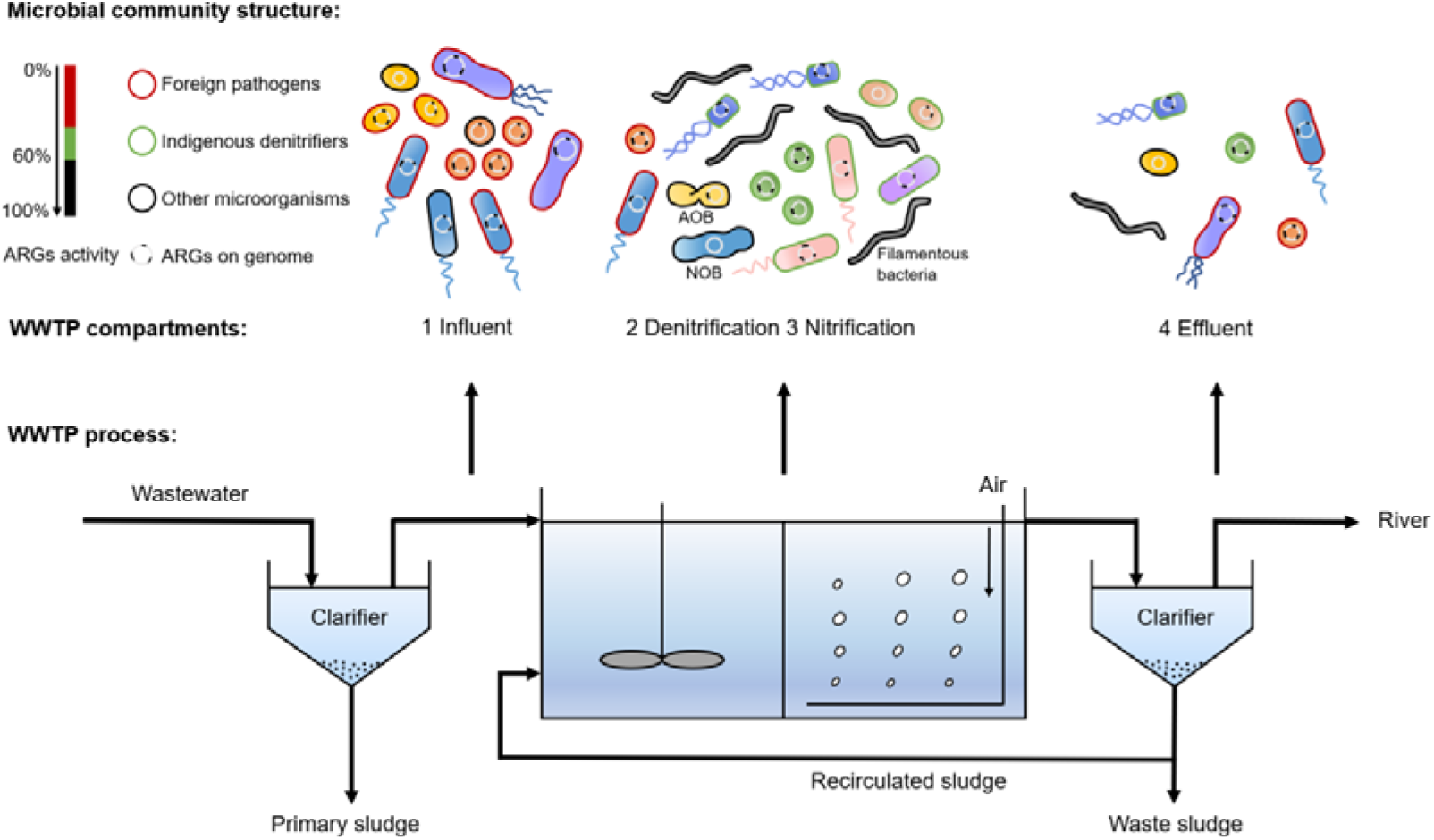
Potential pathogens and indigenous denitrifiers as active and key players of multi-antibiotic resistance in the urban wastewater treatment plants (WWTPs). Microbial samples were taken from the influent, denitrification and nitrification bioreactors, and the effluent of 12 urban WWTP systems. Metagenomic sequencing, assembly and binning together with metatranscriptomic analysis enables a genome-level high-resolution and systematic view on the identity, multi-resistance, pathogenicity, and activity of diverse antibiotic resistance genes (ARGs) hosts throughout the WWTPs. Potential pathogens (marked by red border, defined as MAGs that taxonomically predicted as human pathogens and harbored at least one experimentally verified virulence factor) may derived from human intestinal tracts were abundant in the influent. Diverse microorganisms lived in the denitrifying and nitrifying sludge, including the indigenous denitrifiers (marked by green border, defined as MAGs that shared > 95% total expression activities of denitrification genes in the nitrifying and denitrifying sludge while ≤ 5% total expression activities in the influent and effluent). Most members of potential pathogens and indigenous denitrifiers were identified to host multi-antibiotic resistance genes and were not completely eliminated from the final effluent, thus they represented hitherto-unraveled disseminators of WWTP-released ARGs. Overall, potential pathogens and indigenous denitrifiers contributed ∼60% of all antibiotic resistance activities detected in the recovered genomes and were considered as active and key players of antibiotic resistance in the WWTPs.

The metagenomes and metatranscriptomes generated by our preliminary study have been used to gain an overview on the fate and expression patterns of known antibiotic, biocide and metal resistance genes in the WWTPs^6^. However, the identity, multi-resistance, pathogenicity, distribution, activity and other functional traits of ARG hosts remained unknown, due to the fragmented nature of metagenome assemblies obtained. In this study, we filled the knowledge gaps by re-analysis of the datasets using an advanced genome-centric metatranscriptomic strategy to answer the following questions about bacterial populations hosting ARGs in the 12 urban WWTPs. First, who are the ARG hosts and what are their functional roles throughout the WWTP compartments? Second, which ARGs are likely be mobilized and/or hosted by bacterial pathogens? Third, who are the important ARG hosts that actively express ARGs throughout and across WWTPs especially in the treated effluent? To address these questions, we first resolved the genome phylogenies of active ARG hosts in the WWTPs and found *Proteobacteria* and *Actinobacteriota* as the two most common bacterial hosts. We then checked the multi-resistance, pathogenicity, distribution, activity, survival, and other key functional traits (e.g., biological nitrogen removal) of all the identified ARG hosts from the WWTPs, leading to the key finding that potential pathogens and indigenous denitrifiers are transcriptionally active and key players of wastewater (multi-)antibiotic resistance genes (Fig. 1). This study simultaneously links ARGs to their host identity, activity and functionality in the varying WWTP compartments, which offers a comprehensive, in-depth and new understanding of the key functional traits and microbial ecology of antibiotic resistance in WWTPs.

## MATERIALS AND METHODS

### Genome-centric Reanalysis of WWTPs Microbiome Data

Between March and April 2016, a total of 47 microbial biomass samples were taken from the primarily clarified influent, the denitrifying bioreactors, the nitrifying bioreactors and the secondarily clarified effluent of 12 urban WWTPs that mainly receives domestic sewage across Switzerland. Total DNA and RNA extractions, processing of the mRNA internal standards, data pretreatment, and metagenome assembly were performed as previously described in our earlier publication^6^.

### Genome Binning, Annotation and Phylogenetic Analysis

Metagenome-assembled genomes (MAGs) were recovered using MetaWRAP (v1.2.2)^24^ pipeline. Briefly, with metaBAT2 in the binning module, MAGs were reconstructed from the 47 single-sample assemblies. Contamination and completeness of the recovered MAGs were evaluated by CheckM (v1.0.12)^25^, and only those genomes with quality score (defined as completeness – 5×contamination) ≥ 50 were included in the succeeding analysis. The draft genomes were dereplicated using dRep (v1.4.3)^27^ with default parameters, which resulted in a total of 248 unique and high-quality MAGs. The recovered MAGs were deposited in the China National Gene Bank Database (CNGBdb: https://db.cngb.org/) under the project accession number CNP0001328. The accession numbers of 248 MAGs were listed in Dataset S2.

Taxonomy affiliation of each MAG was determined by GTDB-Tk (v0.3.2)^28^ classify_wf. Open reading frames (ORFs) were predicted from MAGs using Prodigal (v2.6.3)^29^. Phylogenetic analysis of MAGs was conducted with FastTree (v2.1.10)^30^ based on a set of 120 bacterial domain-specific marker genes from GTDB, and the phylogenetic tree was visualized in iTOL^31^.

### ARG Annotation and Mobility Assessment

The annotation of ARGs from the recovered MAGs was accomplished using DeepARG (v2)^32^ with options ‘—align --genes --prob 80 --iden 50’. Predicted ARGs of antibiotic classes with less than 10 reference sequences in the database were removed to avoid mis-annotation due to possible bias. In total, 496 ORFs annotated in 162 MAGs were identified as ARGs with resistance functions to 14 specific antibiotic classes, while 312 ORFs annotated in 117 MAGs were identified as ARGs of multidrug class and were listed in Dataset S3 but not included in the downstream analysis. The 248 high-quality MAGs were then categorized as “multi-resistant” (113), “single-resistant” (49) and “non-resistant” (86), according to whether >1, =1, or =0 ARG classes were annotated in the genome, respectively.

Considering the importance of the plasmid for spreading ARGs, the presence of plasmid sequences in the metagenomic contigs was checked by PlasFlow (v1.1)^33^ which utilizes neural network models trained on full genome and plasmid sequences to predict plasmid sequences from metagenome-assembled contigs. A strict parameter ‘--threshold 0.95’ was employed to robustly compare the occurrence frequencies of plasmid contigs in the binned (i.e., MAGs) and un-binned contigs. Moreover, mobile genetic elements (MGEs) were identified by hmmscan^34^ against Pfam^35^, with options ‘--cut-ga’. The mobility of ARGs was predicted based on either their location on the plasmid contig or co-occurrence with an MGE in a nearby genomic region (<10 kb)^36^.

### Identification of Pathogenic Genomes

The candidate pathogenic genomes were firstly taxonomically identified based on two published reference pathogen lists containing 140 potentially human pathogenic genera^37^ and 538 human pathogenic species^38^. Then, 3642 experimentally verified virulence factors downloaded from pathogenic bacteria virulence factor database (VFDB, last update: Jun 27 2020)^39^ were used to construct a searchable blast database. The ORFs of taxonomically predicted candidate pathogenic genomes were searched against the constructed virulence factor database by BLASTN, and those genomes with an ORF with global nucleic acid identity > 70% to any virulence factor sequence were finalized as belonging to potential human pathogens.

### Nitrification-denitrification Genes Annotation

To explore certain functional traits (i.e., biological nitrogen removal driven by nitrification and denitrification in WWTPs) of ARG hosts in the WWTPs, nitrification-denitrification genes (NDGs) were annotated. Briefly, all MAG-predicted ORFs were searched against a nitrogen cycle database (NCycDB)^40^ using DIAMOND^41^. Those ORFs annotated as nitrification or denitrification genes with global nucleic acid identity > 85% to the reference sequences in the NCycDB database were directly interpreted as functional genes related to nitrogen removal in the WWTPs. Other ORFs were further checked by BLASTN against the NCBI nt database, ORFs with global nucleic acid identity > 70% to the reference sequences were also identified as annotatable functional genes. Together, 283 ORFs from 88 MAGs were annotated as NDGs. With the intention to display the distribution patterns of NDGs in the MAGs, a network was constructed and visualized in Gephi (v0.9.2)^42^. The network was divided into seven parts according to nitrification (3) and denitrification (4) pathway steps.

### Quantitative Analyses of Genome-centric Metatranscriptomics

#### Quantification at the genome level

In order to calculate the relative abundance and the expression level of each MAG, 47 metagenomic datasets of clean DNA reads and 47 metatranscriptomic datasets of clean mRNA reads were mapped across the 47 individual assemblies and the 47 ORF libraries using bowtie2 (v2.3.4.1)^43^, respectively. The resulting .sam files contained mapping information of both MAGs and un-binned contigs, and subsequent filtering extracted mapping results of each MAG. Then, the relative abundance and expression level of each MAG was calculated and normalized to RPKM (reads per kilobase per million) values as the total number of bases (bp) that mapped to the genome, divided by the MAG size (bp) and the sequencing depth (Gb).

#### Quantification at the gene transcript level

To overcome the limitation of relative abundance in the metatranscriptomic analysis^44^, absolute expression values (AEV) were calculated for the 496 ARGs annotated in the 248 high-quality MAGs based on spiked mRNA internal standards^6^ and mapping results. Here, AEV was calculated as ‘transcripts/g-VSS’ (TPG_VSS_) using the following equation:

Absolute expression value (AEV) =

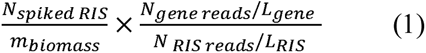

where N_spiked RIS_ is the copy numbers of spiked mRNA internal standards (RIS), *m_biomass_* is the mass of collected volatile suspended solids (VSS) which was regarded as the proxy for biomass by environmental engineers, *N_gene reads_* is the number of reads mapped to the gene in the metatranscriptomic dataset, *L_gene_* is the length of the gene, *N_RIS reads_* is the number of reads mapped to the RIS in the metatranscriptomic dataset, *L_RIS_* is the length of the RIS. This calculation is optimized by weighing different lengths of reported genes, and only genes with >50% of their lengths covered by mapped reads were considered. In this study, if the sample range is not otherwise specified, AEV of a gene refers to the average AEV across all 47 samples.

While AEV is the absolute expression activity of a given gene, relative expression ratio (RER) is a comparison between the given gene and the single-copy marker genes (SCMG) in the genome, which calculated by relativizing the AEV of the given gene by the median AEV of the SCMG in the genome as shown below:

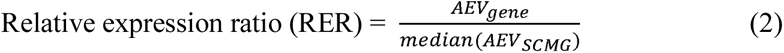

The single-copy marker genes in the recovered genomes were determined by GTDB-tk^28^ which searched 120 ubiquitous single-copy marker genes of bacteria^45^ in the genome, and those unique marker genes in the genome were used to calculate basic expression level of the genome. Ideally, if RER > 1, this gene would be regarded as over expressed compared with the house-keeping marker genes, and if RER = 1, it indicates that this gene expresses at a same level as the marker genes. Similarly, if RER < 1, it indicates that this gene is under expressed compared with the marker genes. Our proposal of these two metrics (i.e., AEV and RER) offer complementary insights into a given gene of interest: AEV quantifies its absolute expression activity in a sample, thus proportionally corresponds to the changing concentration of its host cells within a given microbial community, while RER measures its relative expression compared with basic expression level of its host genome. Thus, RER is a more sensitive parameter to monitor microbial response to environmental changes. Finally, the aggregate AEV and average RER of ARGs in the genome were used to represent the absolute and relative expression activity of the antibiotic resistance function in this genome, respectively.

#### Statistical Analysis

All statistical analyses were considered significant at *p* < 0.05. The similarity of microbial community structure between the nitrification and denitrification bioreactors was examined by mantel test in R using the function ‘mantel’ in the vegan package^46^. The difference of relative expression ratio of individual ARGs and ARGs in the recovered MAGs between the influent and effluent wastewater was determined by Mann-Whitney U test using function ‘wilcox.test’ with option ‘paired=FALSE’ in R. The difference of concentration of antibiotics between the influent and effluent wastewater was determined by Mann-Whitney U test using function ‘wilcox.test’ with option ‘paired=FALSE’ in R. The test of difference in relative expression ratio of ARGs between the four compartments was performed with Kruskal-Wallis test in python using function ‘kruskal wallis’ in scipy package. The average RER of ARGs and denitrification genes in the MAGs was calculated after removing outliers (based on the 3σ principle).

## RESULTS AND DISCUSSION

### Metagenome-assembled Genomes Recovered from the WWTPs Microbiome

The key functions of urban WWTPs such as removal of organic carbon and nutrients are largely driven by uncultured microorganisms^18, 47, 48^. To explore the key microbial functional groups including uncultured representatives, 1844 metagenome-assembled genomes (MAGs) were reconstructed from 47 samples taken from varying compartments in the 12 Swiss WWTPs. A total of 248 unique and high-quality MAGs were retained for further analysis after dereplication and quality filtration. These genomes accounted for 14-62% (average 38%) and 7-75% (average 28%) of paired metagenomic and metatranscriptomic reads, respectively, and therefore represented an important fraction of the microbial community in the WWTPs (Dataset S1). Basic information on the MAGs recovered was listed in Dataset S2. Phylogenetic analysis based on 120 single-copy marker genes of the 248 MAGs showed their grouping and taxonomic classification into 15 phyla (Fig. 2). The MAGs recovered was most taxonomically assigned to *Proteobacteria* (88), followed by *Patescibacteria* (68), *Bacteroidota* (39), *Actinobacteriota* (22), *Firmicutes* (11) and *Myxococcota* (4). The phylum-level microbial community composition in the 12 WWTPs was overall similar to a recent study that recovered thousands of MAGs from activated sludge of global WWTPs that were also mostly assigned to *Proteobacteria*, *Bacteroidota* and *Patescibacteria*^49^.

**Fig. 2.**
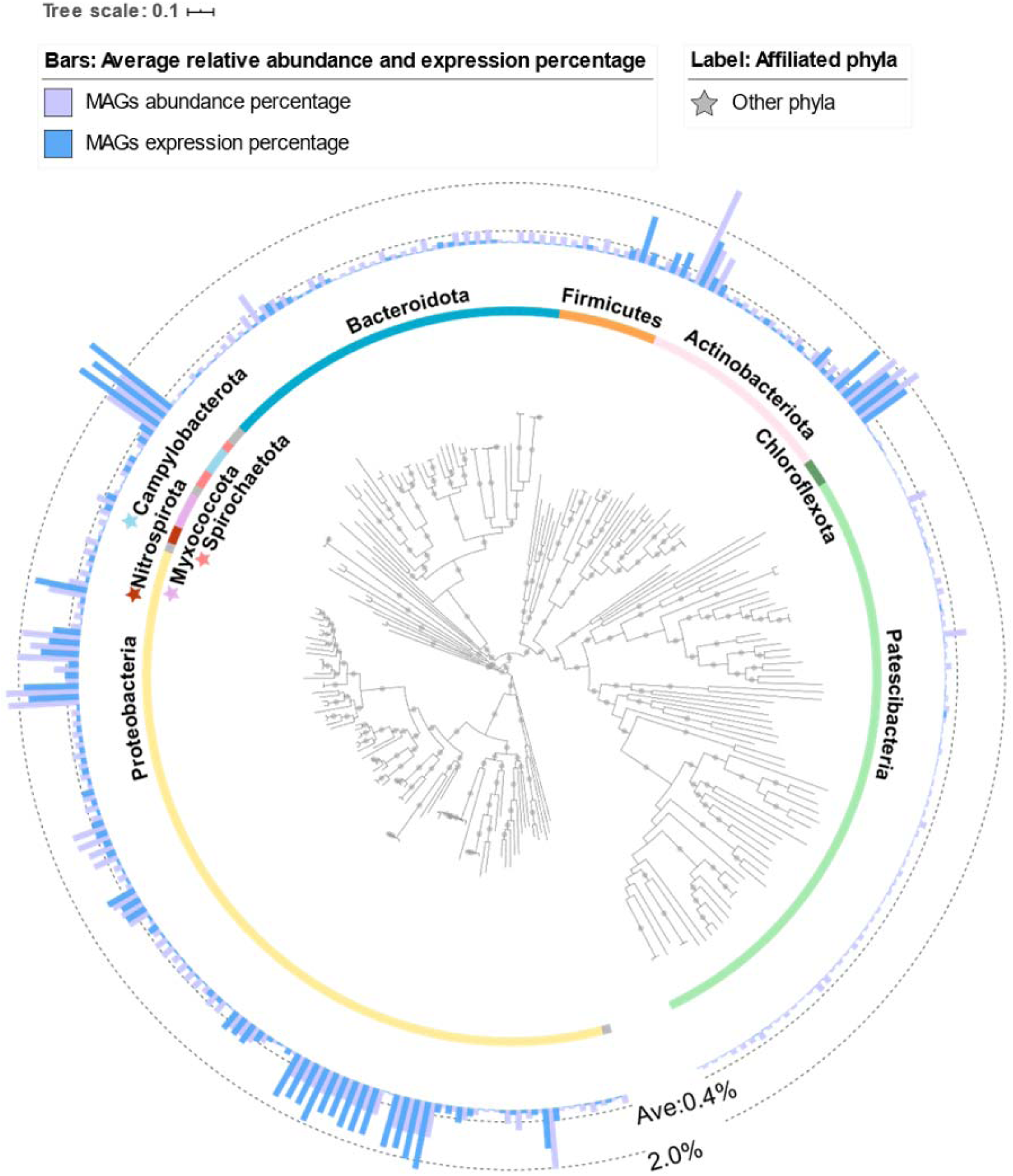
Phylogenetic tree of 248 high-quality MAGs recovered from 12 urban WWTPs. The tree was produced from 120 bacterial domain-specific marker genes from GTDB using FastTree and subsequently visualized in iTOL. Labels indicate phyla names and, to facilitate an easier differentiation, the color of the front stars beside the phyla label is the same as the color of the corresponding phyla; phyla in which only one MAG were recovered were taken as others. The relative abundance and expression level of each MAG were calculated based on RPKM values across all samples. Abundance percentage and expression percentage were proportions of relative abundance and expression level, respectively, and were shown by external bars (purple: abundance percentage; blue: expression percentage). The dashed circles represent the scale for abundance and expression percentage (inside: average 0.4%, outside: 2.0%). Bootstraps >75% are indicated by the grey dots.

Further comparisons of abundance percentage and expression percentage of the 248 MAGs across all samples clearly showed distinct DNA- and mRNA-level compositional profiles across phyla and genomes. Overall, 3, 22 and 88 MAGs assigned to *Campylobacterota, Actinobacteriota* and *Proteobacteria* exhibited a high average abundance percentage of 1.9%, 0.8% and 0.6%, corresponding to an average expression percentage of 4.1%, 0.7%, and 0.7%, respectively. In contrast, *Patescibacteria* showed low average abundance percentage (0.12%) and expression percentage (0.01%). This newly defined superphylum, belonging to a recently discovered candidate phylum radiation^50, 51^, was found to be the second most frequent populations in the 12 WWTPs of this study. These *Patescibacteria* populations, however, might have been overlooked by previous large-scale 16S rRNA-based surveys^17, 47, 52^ due to the special features of their 16S rRNA gene (i.e., encoding proteins and have self-splicing introns rarely found in the 16S rRNA genes of bacteria)^53^. Our first discovery of their survival at extremely low gene expression level (Fig. 2) calls for further investigation of the original sources and potential functional niches of these ultra-small cells (< 0.2 μm) in WWTPs^54^.

### Host Identity, Expression Activities and Mobility of ARGs

To understand taxonomic distribution and activity of ARGs in the MAGs recovered from the WWTPs, a genome-centric metatranscriptomic approach was exploited to examine ARGs in genomic and transcriptomic contexts of all 248 MAGs. Together, 496 ORFs carried by 162 (65.3%) MAGs were identified as ARGs encoding resistance functions of 14 antibiotic classes (Dataset S3). The predicted 162 ARG hosts were further categorized as “multi-resistant” (113 MAGs, 45.6%) and “single-resistant” (49 MAGs, 19.8%) (Fig. 3a, Dataset S2). Among those multi-resistant MAGs, W60_bin3 and W72_bin28 affiliated with *Aeromonas media* and *Streptococcus suis,* respectively, were found to harbor the largest numbers of ARGs, i.e., they both carried 11 ARGs conferring resistance to 9 and 4 antibiotic classes, respectively, followed by 3 MAGs from *Aeromonas media* (2) and *Acinetobacter johnsonii* (1) that carried 10 ARGs (Fig. 3a and Dataset S2).

**Fig. 3.**
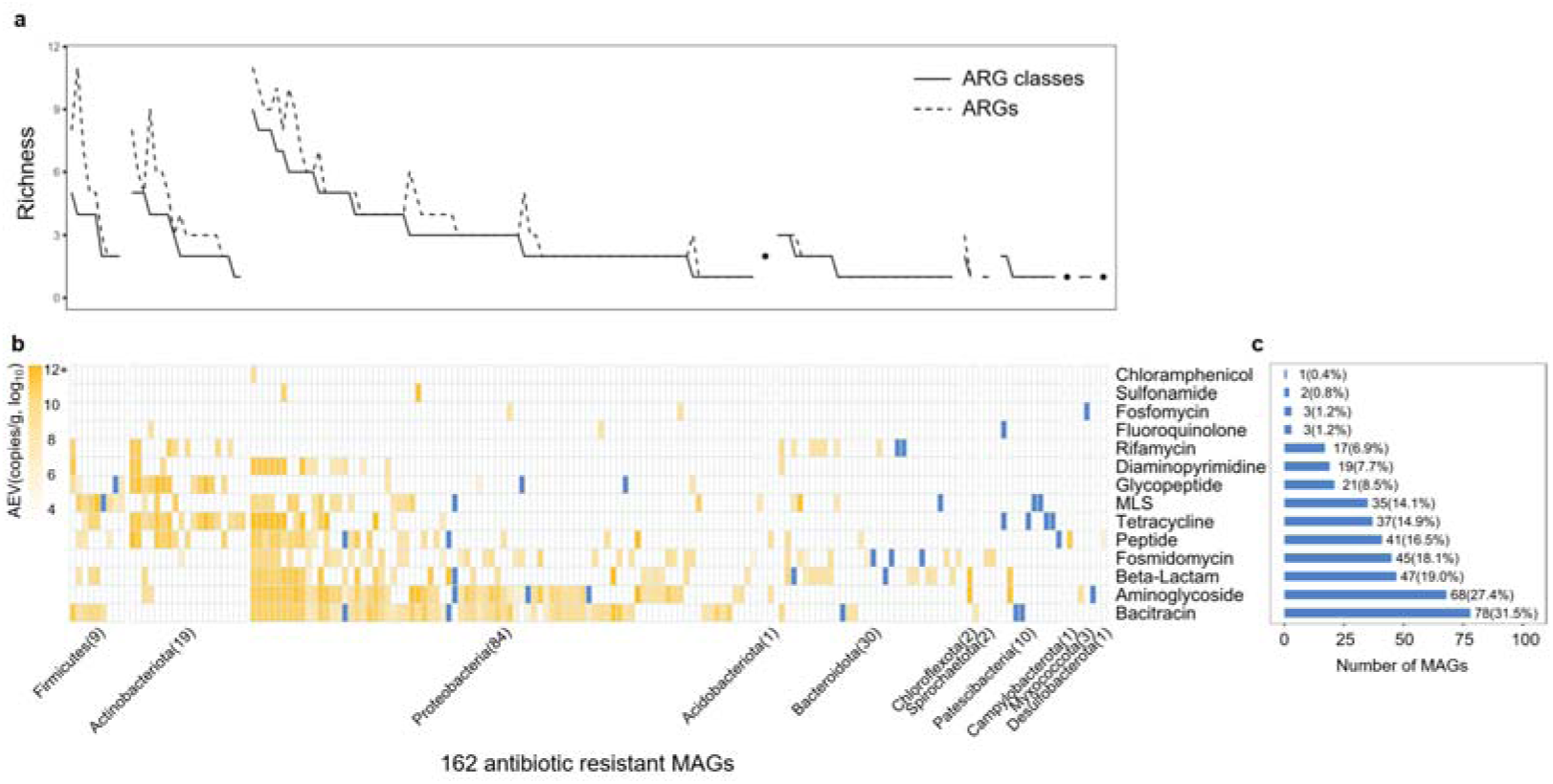
The distribution and activity of ARGs in the recovered genomes. **a.** richness of ARGs and ARG classes detected in 162 resistant MAGs. **b.** taxonomic distribution and absolute expression value (AEV, transcripts/g-VSS) of ARG classes across MAGs. Yellow color intensity represents average AEV of ARGs from each ARG class in the genome. Blue color represents the corresponding MAG harbored but not expressed the corresponding ARG. Figure a and b share the same horizontal axis. **c.** number of MAGs assigned to each class of ARGs.

Taxonomically, ARG hosts were found in 11 out of 15 phyla (except for *Verrucomicrobiota*_A, *Bdellovibrionota*, *Nitrospirota and Gemmatimonadota,* each containing no more than 2 MAGs) (Fig. 3b). MAGs assigned to the phylum of *Proteobacteria* were the most frequent hosts of ARGs. In 88 *Proteobacteria-*affiliated MAGs, 84 MAGs were ARG hosts encoding resistance of 13 antibiotic classes in total, and nearly all of them (83 MAGs) were transcriptionally active for resistance to at least one antibiotic class (Fig. 3b). *Actinobacteriota* were also active hosts of ARGs of 10 antibiotic classes, especially for glycopeptide and tetracycline (Fig. 3b). In contrast, *Patescibacteria* were transcriptionally inactive hosts of ARGs, i.e., 10 out of 68 MAGs encoded ARGs with only one population (W73_bin6) displaying transcription of beta-lactam and aminoglycoside resistance (Fig. 3b). *Patescibacteria* were recently revealed to harbor small but mighty populations with strong adaptability. They usually have reduced genomes (∼1 Mbp) and truncated metabolic pathways^55^, and an under representation of ARGs in their genomes may be a strategic outcome from their process of reducing redundant and nonessential functions.

Among the 14 resistance types of ARGs identified (Fig. 3c), ARGs against bacitracin (78, 31.5%) and aminoglycoside (68, 27.4%), being most prevalent in *Proteobacteria*, were found to be the two most frequent resistance types, followed by ARGs against beta-lactam (47, 19.0%) and fosmidomycin (45, 18.1%). In contrast, sulfonamide- (2, 0.8%) and chloramphenicol-resistance genes (1, 0.4%) were both hosted by few MAGs, all belonging to *Proteobacteria* (Fig. 3b). Absolute quantification revealed that the sulfonamide resistance genes showed the highest expression level with an average AEV of 2.53×10^11^ transcripts/g-VSS, followed by those against tetracycline (1.51×10^11^ transcripts/g-VSS) and peptide (1.46×10^11^ transcripts/g-VSS). In contrast, the fluoroquinolone resistance genes displayed the lowest average AEV (1.42×10^9^ transcripts/g-VSS, Dataset S4). Among all 496 ARGs, 460 ARGs were confirmed to have transcriptional activity in at least one sample (Dataset S4). This indicated that most ARGs are expressed under the environmental condition of the WWTPs. The expression of ARGs could be induced by specific antibiotics or their co-selective or -expressive antimicrobial agents (e.g., other antibiotics and heavy metals) in wastewater, but may also be constitutively expressed or only globally regulated by the metabolic regulators^56^. These results reveal that multiple ARGs were widely distributed and expressed in the WWTPs microbiome.

Plasmids are evolutionarily important reservoir and transfer media for ARGs. From our study, 11 ARGs were found to locate on the plasmid contigs (Dataset S5), three of which were carried by potential pathogens (see Fig. 4) i.e., *tet*39 and ANT(3’’)-IIc carried by *Acinetobacter johnsonii* and *lnu*A carried by *Streptococcus suis* as later discussed. It is notable that plasmid sequences, especially when present in multi-copies or shared across bacteria, are largely excluded from (thus poorly represented) in the reconstructed genomes which are supposed to mainly consist of single-copy genomic regions with nearly the same coverage^57^. For example, our first-hand data from one WWTP showed that only 2.3% contigs from MAGs were predicted by PlasFlow as plasmid sequences, while 6.4%, 7.1%, 7.7% and 9.9% contigs from un-binned contigs assembled from influent, denitrifying sludge, nitrifying sludge and effluent metagenomes were predicted as plasmid-originated. Besides, 35 ARGs identified from the MAGs were located near to a mobile genetic element (MGE, <10kb) including six cases that ARG and MGE are directly adjacent on the same contigs (Dataset S5). These results together reveal possible mobility and thus dissemination potentials of wastewater ARGs mediated by plasmids or other MGEs.

**Fig. 4.**
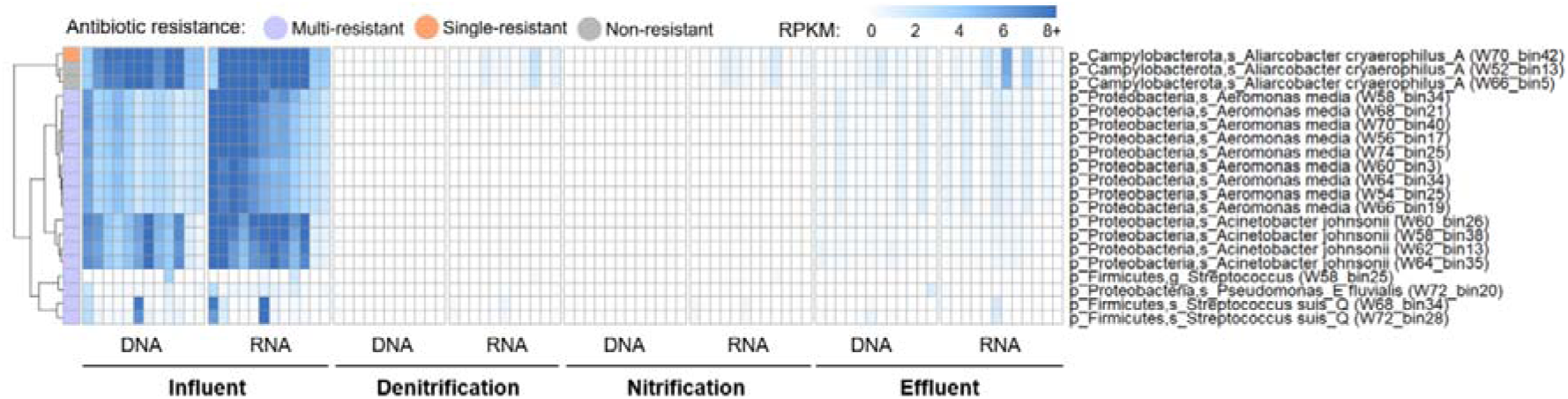
The cross-compartment distribution and expression pattern of potential pathogenic populations in the WWTPs. Heatmap for relative abundance and expression level of 20 potentially pathogenic populations MAGs in the influent, denitrification, nitrification and effluent compartments. Blue color intensity represents genome relative abundance and expression level normalized by RPKM values. Left annotation column shows antibiotic resistant patterns of potential pathogens. Heatmap clustering is computed by “euclidean” distance metric.

### Pathogenicity, Distribution and Activities of ARG Hosts Across WWTP Compartments

Whether environmental ARGs are hosted by clinically relevant pathogens is central to assessing their health risks. Compared with reported metagenomic contigs or gene fragments^58, 59^, MAGs provide a more complete genome context allowing for more robust host identification at higher resolution, down to the genus or species level. In total, 20 potentially pathogenic MAGs were identified based on the published reference pathogen lists^37, 38^ and verified the presence of virulence factors. Seventeen out of the 20 pathogenic MAGs were found to encode multi-antibiotic resistance, and the aforementioned 5 MAGs that encode the largest number of ARGs (10 or 11) all belonged to the pathogenic group. The potentially pathogenic organisms overall accounted for 47.3% abundance and 65.4% expression activity in the influent samples (Dataset S6). These potentially pathogenic bacteria were abundant and active in the influent sewage and likely originated from the human intestinal tract. It is noteworthy that members of pathogenic group were almost absent in the downstream denitrifying and nitrifying bioreactors but were observed again in the effluent where they were not completely eliminated (Fig. 4). We suspected that these influent-abundant pathogens were mainly planktonic cells that generally failed to invade or inhabit activated sludge flocs, but passively drifted into the final effluent with wastewater flow. Among the 20 pathogenic MAGs, 3 were assigned to *Aliarcobacter cryaerophilus*, a globally emerging foodborne and zoonotic pathogen which may cause diarrhea, fever, and abdominal pain to human^60^. *A. cryaerophilus* showed high abundance and expression activity in the influent samples (Fig. 4) and they were confirmed to present in food of animal origin, drinking water, and sewage before^61^. Although these three *A. cryaerophilus* species were classified as either non-resistant or single-resistant, their considerable transcriptional activities in the effluent (average RPKM in effluent > 1, Dataset S6) deserve further attention. The 9 MAGs classified as *Aeromonas media*, a well-known gram-negative, rod-shaped and facultative anaerobic opportunistic human pathogen^62^, were all identified as being resistant to more than three classes of antibiotics and transcriptionally active in the effluent (RPKM in effluent: 0.58∼0.67, Dataset S6). In addition, other potential pathogens survived wastewater treatment included *Acinetobacter johnsonii* (4 MAGs), *Streptococcus* (3 MAGs) and *Pseudomonas fluvialis* (1 MAG) (Fig. 4 and Dataset S6). Together, 18 antibiotic resistant pathogens from the wastewater influent may have roles as persistent pathogenic agents and ARGs disseminators in the WWTP effluents, as they could successfully enter into the receiving rivers where health risks associated with their local propagation, resistance transfer and human exposure call for research attention.

The comparative profiles in relative abundance and expression level of the 162 ARG hosts as well as the 86 non-resistant MAGs across 47 samples showed that both the population distribution and the expression profiles dramatically shifted across influent, denitrification, nitrification, and effluent compartments (Fig. S1), probably driven by environmental heterogeneity and habitat filtering. Interestingly, although the denitrification and nitrification compartments differed significantly (paired t-test *p*< 0.001) in dissolved oxygen (0.02±0.004 vs. 2.04±0.17 mg/L), organic carbon (14.32±1.48 vs. 11.23±1.42 mg/L), ammonia nitrogen (8.06±1.02 vs. 2.26±0.63 mg/L), nitrate nitrogen (3.94±1.24 vs. 8.82±1.33 mg/L) and hydrolytic retention time (3.92±0.45 vs. 8.33±1.30 days)^6^, the two compartments shared almost the same genomic and transcriptomic composition (mantel statistic r = 0.900 and 0.957, *p*< 0.001; Fig. 4), suggesting that a set of core species can survive and thrive in the classic anoxic-aerobic cycles of activated sludge process. Unlike the tightly clustered profiles in the influent, the effluent had highly dispersive population distribution and expression patterns that partially resembled those of activated sludge and influent, revealing prominent impacts from wastewater treatment and diverse emission of viable resistant bacteria.

### Multi-antibiotic Resistance Associated with Biological Nitrogen Removal

Biological nitrogen removal is one of the key goals of wastewater treatment processes. It is driven by nitrifiers and denitrifiers which were found to be closely associated with antibiotic resistance in this study. Together, 88 MAGs were found to be potentially involved in wastewater nitrogen removal (Dataset S2). Compared with nitrification, a much higher diversity of microbes (7 phyla vs. 3 phyla, 87 vs. 5 unique MAGs) showed genetic potential for denitrification. There were 8 MAGs from *Proteobacteria* expressed genes for full denitrification (i.e., NO_3_ - NO_2_ - NO - N_2_O - N_2_) and other 79 MAGs expressed genes for partial denitrification. This finding from WWTP systems echoed the widely accepted ecological concepts that nitrification is often carried out by specialist taxa while denitrification can involve a wide range of taxa^63^. It was noteworthy that 4 MAGs simultaneously expressed denitrification and nitrification genes (2 MAGs from *Nitrospira*, 1 MAG from *Nitrosomonas* and 1 MAG from *Caldilineales*, Dataset S2). Detailed description of nitrification-denitrification genes (NDGs) distribution in the 88 MAGs is available in the Supplementary Information S1, suggesting the presence of these functional bacteria and genes as the basis for biological nitrogen removal from wastewater.

Among these MAGs, a portion of nitrifying populations (3/5 MAGs) and most of denitrifying (without nitrifying) populations (75/83 MAGs) were multi-resistant (71/88 MAGs) or single-resistant (7/88 MAGs), while the majority of non-resistant populations (76/86 MAGs) were not involved in either nitrification or denitrification (Fig. 5b), revealing antibiotic resistance maybe an important trait for successful survival and routine functioning of nitrogen-removing bacteria under WWTP conditions, i.e., in the presence of wastewater-borne antimicrobial stressors. The two ammonia-oxidizing MAGs classified as *Nitrosomonas* (W68_bin8 and W79_bin32), both expressed ARGs of bacitracin, and W68_bin8 additionally expressed ARGs of fosmidomycin and tetracycline. The two nitrite-oxidizing MAGs classified as *Nitrospirota* (W81_bin21 and W77_bin34) did not encode detectable ARGs. Besides, 306 out of 496 ARGs were in the MAGs of potential denitrifiers, revealing that denitrifying bacteria are important hosts of diverse ARGs in the WWTPs (Dataset S2). The high prevalence of ARGs in denitrifiers was reasonable because there was some evidence showing the existence of antibiotics would cause a significant inhibition to denitrification genes^64–66^. Considering the presence of various antibiotics in the WWTPs (Dataset S8), denitrifiers carrying ARGs could better maintain their denitrifying function and protect themselves from inhibition by the antibiotics. When both taxonomic affiliation and nitrogen removal function of the 248 MAGs were considered, we found that multi-antibiotic resistant *Proteobacteria* (58/88 MAGs, 65.9%) played a predominant role in the nitrification and denitrification, while *Patescibacteria* (66/68 MAGs, 97.1%) and *Bacteroidota* (26/39 MAGs, 66.7%) were dominated by non-resistant or single-resistant populations without a detectable NDG (Dataset S2). Combined, the above results strongly indicate the high prevalence of ARGs in nitrogen-removing functional organisms, especially denitrifying *Proteobacteria*, a hotspot of multi-antibiotic resistance in WWTP systems. If ARGs are widely distributed in microbes that performing a central function of the WWTP process, they can thus likely not be easily removed from these systems.

**Fig. 5.**
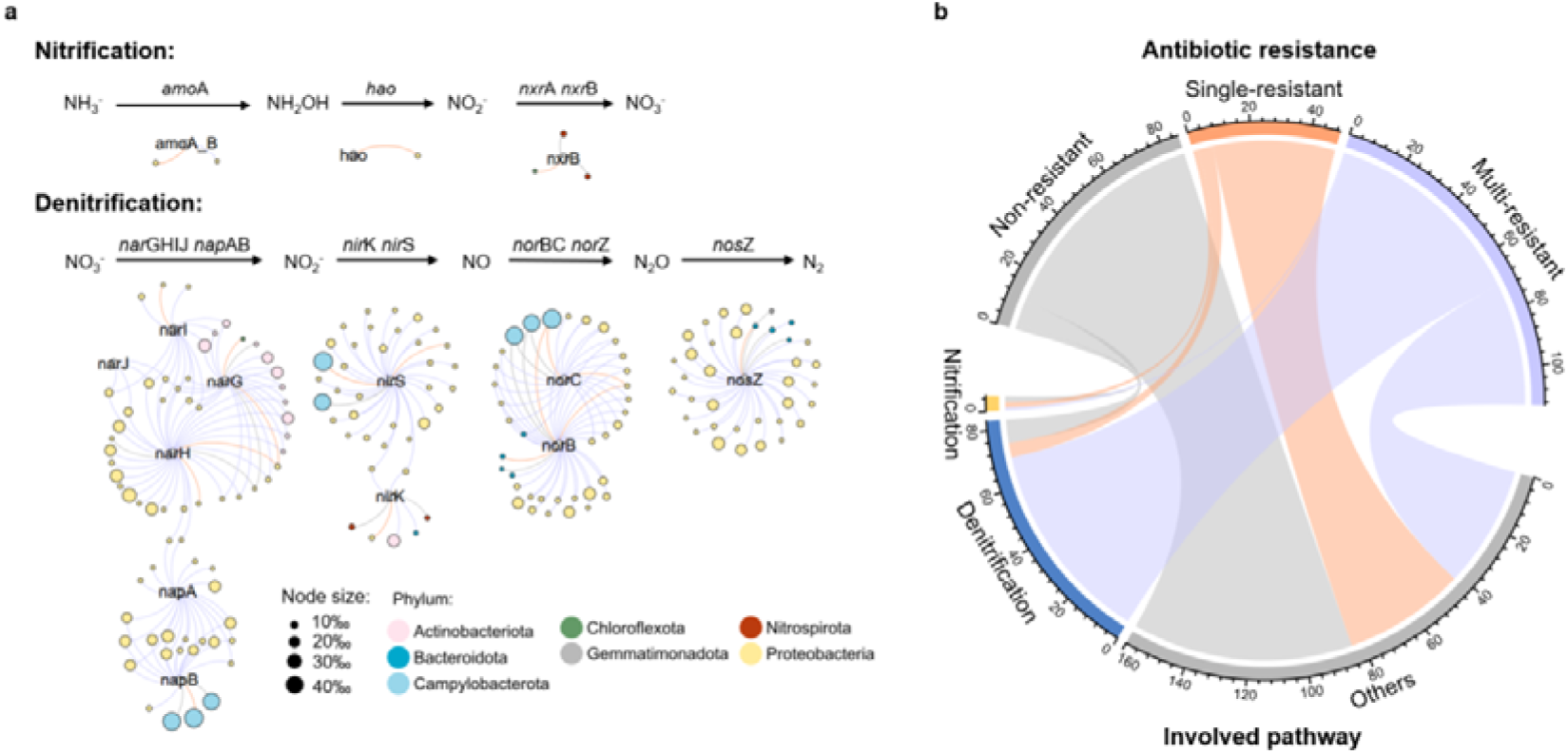
The distribution of MAGs annotated with NDGs and their relationship with antibiotic resistance. **a.** Network reveals distribution of NDGs in nitrifiers and denitrifiers. Each node represented a NDG or MAG (colored by taxonomy and size scaled by expression percentage), and each edge connected a MAG to a NDG which represented the MAG expressed the NDG in at least one sample. Color of edge represents antibiotic resistant pattern of the linked MAG (purple: multi-resistant, orange: single-resistant, grey: non-resistant) **b.** Relationship between antibiotic resistance and nitrogen-removing metabolism in the related MAGs. The width of the string represents the number of MAGs.

### Differential Antibiotic-Resistant Activities across WWTP Compartments

The absolute expression and relative expression levels of ARGs were examined both in the functional groups involved in nitrogen removal (Fig. 6a) and other resistant members (Fig. 6b) across WWTP compartments. Notably, 14 out of 18 resistant pathogens were also identified as denitrifiers, thus they may participate in biological nitrogen removal from wastewater (Fig. 6a). The 18 resistant pathogenic populations (e.g., MAGs from *Acinetobacter johnsonii* and *Aeromonas media*) were found actively expressing ARGs in the WWTPs, and they overall contributed to ∼38% of ARGs expression in the recovered MAGs (Fig. 6a and Dataset S7). Although nitrifiers were overall not active in the expression of ARGs (e.g., W68_bin8 from *Nitrosomonas*: 2.04×10^8^ transcripts/g-VSS, W68_bin12 from *Caldilineales*: 4.03×10^8^ transcripts/g-VSS), some denitrifiers, especially those indigenous denitrifiers (shared >95% total activities of denitrification genes in the nitrifying and denitrifying sludge, ≤5% total activities in the influent and effluent) highly expressed ARGs in the WWTPs (e.g., 3 MAGs from *Phycicoccus* and 2 MAGs from *Tetrasphaera* > 6×10^11^ transcripts/g-VSS). This contrasting pattern between nitrifying and denitrifying bacteria suggests considerable differences in their resistance response and survival strategy to tackle the stresses of antibiotics (Dataset S8) or co-selective antimicrobial agents in the wastewater. Together, the resistant members from potential pathogenic group (marked in red, Fig. 6) and indigenous denitrifying group (marked in green, Fig. 6a) contributed to ∼60% of ARGs expression in the recovered MAGs (Dataset S7). They were both key hosts of ARGs actively expressing ARGs in the WWTPs.

**Fig. 6.**
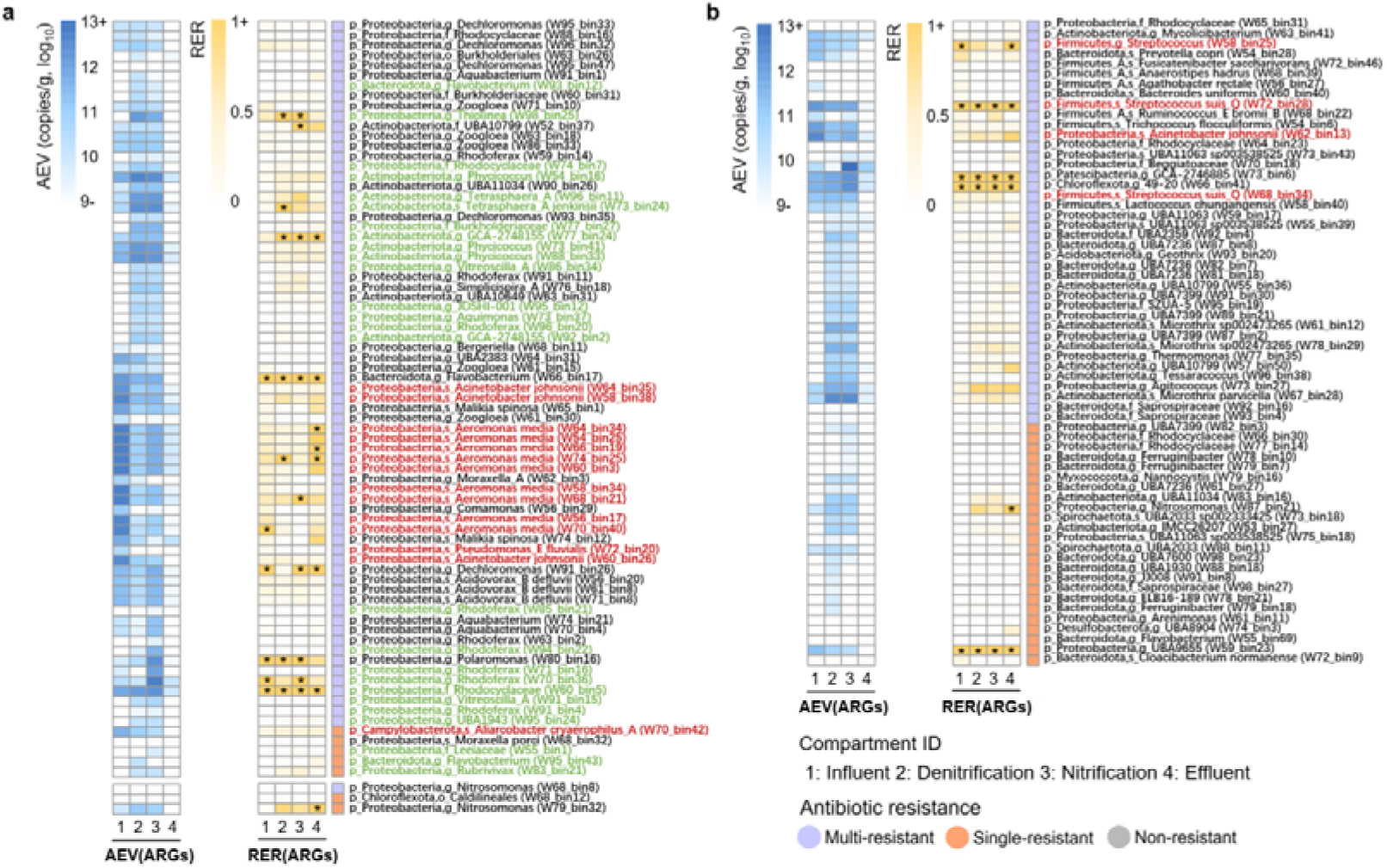
The absolute expression value (AEV) and relative expression ratio (RER) for ARGs in the WWTP bacterial populations. **a.** the AEV and RER of ARGs in MAGs putatively involved in nitrogen removal. **b.** the AEV and RER of ARGs in other MAGs. The case of RER>1 is marked with an asterisk. Right annotation column illustrated antibiotic resistant pattern of MAGs. Column names of heatmap represent compartment ID in the WWTPs. MAGs marked in red were potentially pathogenic group and MAGs marked in green were indigenous denitrifying group.

Of the 64 resistant MAGs without an identifiable NDG but expressed ARGs in the WWTPs, 35 MAGs primarily expressed ARGs in the nitrifying and denitrifying bioreactors (>95% total activities) rather than in the influent and effluent (≤5% total activities, Fig. 6b, Dataset S7). These indigenous resistant bacteria of activated sludge were dominated by populations of phylum *Bacteroidota* (15 MAGs), *Proteobacteria* (11 MAGs) and *Actinobacteriota* (8 MAGs, Fig. 6b). For instance, chemoorganotrophic *Microthrix* (3 MAGs) are associated with activated sludge flocs formation and filamentous bulking^67^, while chemolithoautotrophic *Gallionellaceae* (4 MAG assigned to UBA7399), a poorly characterized family in WWTPs microbiome, are known to harbor aerobic nitrite-oxidizing bacteria (e.g., *Nitrotoga*^68^) and ferrous iron-oxidizing bacteria^69^. Unsurprisingly, the absolute expression of ARGs decreased dramatically (>99%) in most effluent populations, due to the efficient removal of bacterial cells in the WWTPs (e.g., 88%-99%^6^). However, the effluent had witnessed detectable expression of ARGs in the 121 resistant MAGs (Dataset S7). There were 6 multi-resistant MAGs maintained high absolute expression (AEV > 1×10^10^ transcripts/g-VSS) in the effluent, among which, two denitrifying *Malikia spinosa* strains (Fig. 6a) and one *Beggiatoaceae* spp. (Fig. 6b) were identified as the three most pronounced contributors of multi-antibiotic resistant activities in the effluent microbiota (8.88×10^10^, 2.56×10^10^ and 1.67×10^10^ transcripts/g-VSS, respectively). Besides, according to the measurement data of antibiotics in the previous publication^6^, several kinds of antibiotics (e.g., macrolides, clindamycin, vancomycin) were not eliminated significantly (Dataset S8). These residual pharmaceuticals and surviving antibiotic resistant bacteria entering into the receiving water environment may promote the emergence and transmission of ARGs.

While the comparative profiles of absolute expression (i.e., AEV dynamics) enable us to sort out host bacteria actively expressing ARGs, RER provides an additional insight into the relative expression and regulation of ARGs under varying wastewater stresses and environmental changes throughout WWTPs. Overall, relative expression of ARGs were only ∼0.4-fold of the average expression level of the single-copy genes in the host genomes, implying that antibiotic resistance was a generally inactive function with below-average expression level in the WWTP microbiome. Moreover, most ARGs exhibited relatively stable RER dynamics across compartments (Fig. 6). Of 130 MAGs that expressed ARGs in the influent and/or effluent, only 16 (e.g. 2 MAGs from *Zoogloea*) showed significant decrease (Mann-Whitney FDR-*p*< 0.05) in the RER of ARGs from influent to effluent, and 27 (e.g. 5, 3, 3, 2 MAGs from *Aeromonas media*, *Acinetobacter johnsonii*, *Phycicoccus* and *Nitrosomonas*, respectively) showed significant increase (Mann-Whitney FDR-*p*< 0.05) in the RER of ARGs. In contrast, no significant change was observed for the remaining majority MAGs (87/130, 66.9%) (Dataset S9). This result was consistent with the observation at the level of ARGs (Dataset S10, Supplementary information S2), indicating that the transcription of ARGs was overall weakly affected by changing environmental conditions within WWTPs. However, the expression pattern of NDGs was quite different from that of ARGs. The relative expression of denitrification genes and nitrification genes were 4.6-fold and 80.1-fold of average level in the host genomes, respectively, indicating that biological nitrogen removal is a functionally important and metabolically active bioprocess in the WWTPs. The significant upregulation (Mann-Whitney FDR-*p*< 0.05) of denitrification genes from influent to the downstream activated sludge bioreactors was noted in ∼53% of the denitrifiers (46/87 MAGs) (Dataset S9). Notably, two multi-resistant denitrifying populations assigned to *Rhodocyclaceae* and *Flavobacterium* (Fig. 6a), together with four functionally unassigned populations associated with *Streptococcus*, *GCA-2746885*, *49-20* and *UBA9655* (Fig. 6b), actively expressed ARGs (RER >1) across the four treatment compartments. Therefore, these persistently active resistant populations were important reservoirs of wastewater-borne antibiotic resistance.

### Research Significance and Methodological Remarks

To the best of our knowledge, this is the first study to gain so far the most complete insights into the key functional traits of ARG hosts in WWTPs based on both absolute expression activity of ARGs and their relative expression activity in the host genomes. Our findings demonstrated that potential pathogens and indigenous activated sludge denitrifiers in the WWTPs were important living hosts and hotspots of ARGs in which multi-antibiotic resistance genes were not only present but also expressed even in the treated effluent. Further, the almost unchanged relative expression of ARGs in most resistant populations and those resistant bacteria surviving wastewater treatment indicate that these populations are robust under environmental conditions and leave the WWTPs alive, raising environmental concerns regarding their role in dissemination of multi-antibiotic resistance into downstream aquatic ecosystems. Future studies are thus needed to examine the propagation and health risks of wastewater-derived multi-antibiotic resistance determinants with regards to their ability to successfully colonize the receiving environment of and/or regarding human exposure to their pathogenic hosts via such environmental reservoirs.

Our study also demonstrates a new methodological framework that integrates metagenome-centric genomic and quantitative metatranscriptomic analyses to overcome the limitations of existing DNA read-based, gene-based and/or contig-based metagenomic approaches commonly employed for host tracking and risk assessment of environmental ARGs: (i) poor taxonomic resolution, (ii) lack of resistance activity monitoring, and (iii) lack of absolute quantification of ARGs. This new meta-omics framework is not only directly applicable for host tracking of ARGs in other environmental samples or of functional genes other than ARGs, but also sets a foundation for developing related bioinformatics pipelines and tools. Despite the demonstrated power of the framework in resolving key host traits of ARGs, its metagenome-assembled genome analysis necessarily focused on chromosomal ARGs while underestimated plasmid ARGs, although we also recovered resistance contigs of plasmid origin from the MAGs recovered (Dataset S5). Notably, it is hard to link (mobile) multi-resistance plasmids with their host phylogeny with the same confidence as for chromosomal MAGs, nor can it be completely excluded that the bacteria from the studied MAGs do not harbor additional ARG on plasmids, whether an ARG can be identified on their host chromosomes. As the importance of plasmids for spreading antibiotic resistance is well known, the current approach cannot capture the full picture of ARG-host relationships. This limitation of our study would, at least in theory, be circumventable by a massive application of single-cell genomics although at present this approach would still be limited in practice by cost and labor considerations. On the other hand, this study focused on gene activity at the transcriptional level, but lack of information about the actual translated protein. Further metaproteomics study can help to overcome the loss of information about protein, but the potential of the function (i.e., antibiotic resistance in the WWTPs) still needs to be emphasized.

## ASSOCIATED CONTENT

### Supporting Information

Distribution and expression activity of NDGs in functional MAGs involved in nitrogen removal in the WWTPs; Statistical analysis for individual ARGs; Figure showing the cross-compartment distribution and gene expression pattern of all 248 bacterial populations in WWTPs (DOCX)

Datasets showing the percentage of metagenomic and metatranscriptomic reads mapped to the 248 MAGs recovered; genome statistics and genes annotation of 248 MAGs recovered from 12 WWTPs; the annotation results of ARGs from recovered MAGs based on DeepARG; absolute expression value (AEV, transcripts/g-VSS) and relative expression rate (RER) of 496 ARGs in 162 resistant MAGs across 47 samples; co-occurrence instances of ARGs and MGEs on the same resistance contigs from recovered MAGs; relative abundance and expression level of 248 MAGs (as RPKM) in the WWTPs metagenomes and metatranscriptomes; the average expression value (AEV) and relative expression ratio (RER) of ARGs in the MAGs; mann-Whitney test for relative expression ratio (RER) of ARGs and NDGs in the recovered MAGs; Mann-Whitney and Kruskal-Wallis test for the relative expression ratio (RER) of ARGs (XLSX)

## AUTHOR INFORMATION

### Authors Contributions

F. J. designed the experiments and F. J. and L. Y. wrote the manuscript. L. Y., Y. W, and A. P. performed the bioinformatics analysis. L. Y. and L. Z. performed the statistical analysis. J. Z., B. S. and B. H. provided constructive suggestions to the analyses and revised the manuscript. F. J. supervised the project.

### Notes

The authors declare no conflict of interest.

## ACKNOWLEDGMENTS

We thank Dr. WeiZhi Song at the UNSW Sydney and Mr. Guoqing Zhang at the Westlake University for helpful discussion on the part of the bioinformatics procedures.

## DATA AVAILABILITY

The sequence datasets are deposited in China National GeneBank (CNGB) with an accession number CNP0001328.

