## Supplementary material for "Pathogenic and Indigenous Denitrifying Bacteria are Transcriptionally Active and Key Multi-Antibiotic Resistant Players in Wastewater Treatment Plants": SI

**Legends**

**Supplementary Information**

**Supplementary Text S1:** Distribution and expression activity of NDGs in functional MAGs involved in nitrogen removal in the WWTPs

**Supplementary Text S2:** Statistical analysis for individual ARGs

**Supplementary Figure**

**Figure S1:** The cross-compartment distribution and gene expression pattern of all 248 bacterial populations in WWTPs.

**Supplementary Information S1: Distribution and expression activity of NDGs in functional MAGs involved in nitrogen removal in the WWTPs**

Together, there were 283 ORFs from 88 MAGs annotated as nitrification-denitrification genes (NDGs). The distribution, absolute expression value (AEV) and relative expression rate (RER) of NDGs could be briefly described as below.

**Nitrification** Generally, 5 MAGs from 3 phyla encoded functional genes involved in nitrification in WWTPs. The expression profiles of these functional genes showed that ammonia-oxidizing bacteria (AOB) closely collaborated with nitrite-oxidizing bacteria (NOB) to catalyze wastewater ammonia conversion into nitrate in two steps, suggesting the absence of complete ammonia oxidizers (i.e., comammox[1]) in the full-scale WWTPs examined. Specifically, 2 *Nitrosomonas* MAGs of phylum *Proteobacteria* (W68_bin8 and W79_bin32) first oxidized ammonia to hydroxylamine with ammonia monooxygenase subunit A (*amo*A). In contrast to the low AEV of *amo*A in W68_bin8 (2.47$\times$10^9^ copies/g), W79_bin32 represented a transcriptionally active ammonia oxidizer in all WWTP samples at thousands of times higher expression levels (7.34$\times$10^12^ copies/g), revealing its strong activities and thus adaptability in WWTPs. This MAG of *Nitrosomonas* was the only identifiable host of hydroxylamine dehydrogenase (*hao*, 2.25$\times$10^12^ copies/g) for hydroxylamine oxidization into nitrite. Its notable ownership of three denitrification genes including *nirK*, *norB* and *norC* (Dataset S2) suggests that this denitrifying ammonia oxidizer can potentially convert self-produced nitrite into nitrous oxide. Upon ammonia oxidation to nitrite, two *Nitrospira* MAGs (W77_bin34 and W81_bin21) further oxidized nitrite to nitrate. Their nitrite oxidoreductase beta subunit (*nxr*B) showed high AEV of 7.69$\times$10^12^ and 4.91$\times$10^12^ copies/g, respectively. Surprisingly, we found that W68_bin12 belonging to a yet-to-be-cultured species of order *Caldilineales* (Dataset S2), also carried *nxr*B gene, though only slightly expressed in one nitrifying activated sludge sample (1.25$\times$10^9^ copies/g), revealing potential activities in nitrite oxidation. Nitrite oxidizers have been rarely reported in the phylum *Chloroflexi*[2]*.* Whether the *nxrB* gene in this MAG corresponds to the nitrite oxidizing activity of *Chloroflexi* warrants further verification.

**Denitrification** In total, 87 MAGs from 7 phyla were potentially involved in the reduction of nitrate to nitrogen gas. First, 43 MAGs encoding nitrate reductase (*nar*GHIJ operon), together with 25 MAGs encoding periplasmic nitrate reductase (*nap*A) or cytochrome c-type protein (*nap*B), had the capacity to reduce nitrate to nitrite. Then, nitrite was further reduced to nitric oxide by 38 MAGs which expressed nitrite reductase *nir*S and/or *nir*K genes. 37 MAGs encoding nitric oxide reductase (*nor*BC) had the potential to reduce nitric oxide to nitrous oxide, while another type of nitric oxide reductase (*nor*Z) was not detected in the recovered genomes. Finally, 36 MAGs encoded nitrous oxide reductase (*nos*Z) which reduces the nitrous oxide to nitrogen gas (Fig. 4a). The aforementioned MAGs involved in the denitrification processes were dominantly assigned to *Proteobacteria* (64 MAGs including 61 from *Gammaproteobacteria* class and 43 from *Burkholderiales* order) and *Actinobacteriota* (12 MAGs). Of these MAGs, 8 MAGs from *Proteobacteria* (5 from *Rhodocyclaceae*, 2 from *Burkholderiaceae* and 1 from *Pseudomonadaceae*) expressed denitrification genes for the four steps of the complete pathway; 8 MAGs from *Campylobacterota* or *Proteobacteria* expressed the denitrification genes for the first three steps, and 25 MAGs from *Actinobacteriota* or *Proteobacteria* only expressed the denitrification genes for the first step, nitrate reduction (Dataset S2). In aggregate, we observed a decreasing number of MAGs involved in each of the four denitrification steps, in order from first to last step: 66 > 38 > 37 > 36 MAGs. This niche partitioning between wastewater denitrification tasks indicates that metabolic imbalance and the resulting accumulation of intermediates (e.g. potent greenhouse gas nitrous oxide[3]) may considerably occur in WWTPs. Combined, the organic integration and balance of these NDGs and metabolic pathways formed the basis for engineering nitrogen cycle for nitrogen decontamination in WWTP systems in wastewater.

**Supplementary Information S2: Statistical analysis for individual ARGs**

Among 424 ARGs that expressed in the influent and (or) effluent, only 42 ARGs showed significantly decreased RER from influent to effluent (Mann-Whitney FDR-*p* < 0.05), while 52 ARGs showed significantly increased RER (Mann-Whitney FDR-*p* < 0.05). In contrast, no significant change was observed for the remaining majority (330/424, 77.8%) (Dataset S10). Furthermore, Kruskal-Wallis test was used to verify whether ARG expression was significantly different in the four compartments and showed that 341 out of 460 (74.1%) expressed ARG showed no significant change between four compartments (Dataset S10), leading to the same hypothesis that ARGs are overall weakly affected by changing environmental conditions within WWTPs.

**Figure S1: The cross-compartment distribution and gene expression pattern of all 248 bacterial populations in the wastewater treatment plants.** RPKM: reads per kilobase million. Heatmap for DNA-level and RNA-level relative abundance and expression level of metagenome-assembled genomes (MAGs) recovered in the influent, dentification, nitrification and effluent metagenome (i.e., DNA) and metatranscriptome (i.e., RNA), respectively. The cell color intensity represents genome relative abundance and expression level normalized by RPKM values. Left annotation column shows antibiotic resistant patterns of pathogens. The recovered 248 MAGs were distributed unevenly in each sample. In order to make it easy to differentiate the color of the heatmap, the abundance and expression level of MAGs are visualized within the range of 0-2 (RPKM), but the quantification of a few numbers of high-abundance/activity MAGs are unable to distinguish (e.g., max abundance of MAGs in the samples: RPKM 90.34; max expression level of MAGs in the samples: RPKM 99.90, Dataset S6).

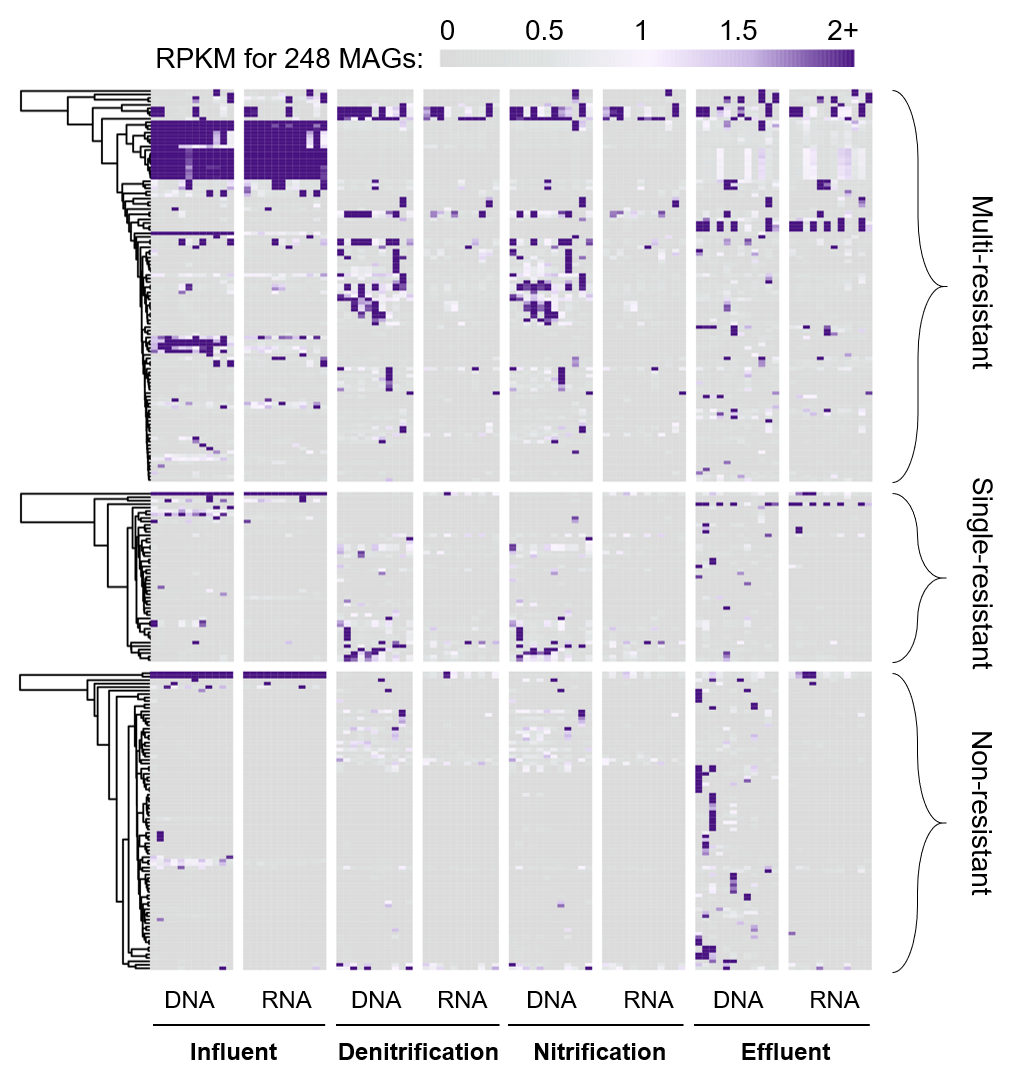
